# Spontaneous activity synchronizes whisker-related sensorimotor networks prior to their maturation in the developing rat cortex

**DOI:** 10.1101/176800

**Authors:** David A. McVea, Timothy H. Murphy, Majid H. Mohajerani

## Abstract

A prominent feature of the cortical systems controlling the whiskers in the adult rodent is tight coupling between sensory and motor systems. Stimulation of the whiskers evokes activation of discrete motor regions of cortex shortly after activation of the sensory cortex. To explore the factors that direct the development of sensorimotor functional connectivity, we recorded spontaneous and whisker-evoked cortical activity using voltage-sensitive imaging over a large (7X7 mm) craniotomy in postnatal rats (day 5-12) under anesthesia. We found that spontaneous bursts of activity in the barrel cortex were correlated predominantly with activity in motor (anterio-medial) cortex, at ages before whisker stimulation evoked activation in this area. Intracortical microstimulation and anatomical tracing experiments confirmed there were no functional or anatomical intracortical sensorimotor connections. We interpret these results as evidence that the spontaneous patterns of activity in the cortex synchronize functionally related regions of the brain prior to their maturation.

**Author Contributions:** D.A.M, T.H.M., and M.H.M. designed the study. D.A.M, and M.H.M performed the experiments and analyzed the data and wrote the manuscript, which all authors commented on and edited. T.H.M. and M.H.M. supervised the study.

---

Integration of sensory and motor signals is necessary to accurately sense the environment and move within it ^1^. A clear example of such integration exists in the whisker sensorimotor system of the rodent ^2^. The barrel cortex, the primary sensory destination of whisker related sensory inputs, is connected to the motor cortex via mono-synaptic excitatory fibers ^3, 4, 5^ and stimulation of the vibrissae or their afferents evokes activity in the motor cortex shortly after activation of the primary barrel sensory cortex ^4, 6, 7^.

How the cortico-cortical connections that underlie these interactions develop is not well understood. Molecular cues under the control of regional gene expression play important roles in the guidance of thalamocortical axons and the subsequent development of cortical identity ^8, 9^ and are likely to have similar roles in the guidance of cortico-cortical neurons ^9, 10^. The contribution of extrinsic, activity dependent factors in the development of cortical sensorimotor circuits is not well known, but activity-independent factors alone are insufficient to allow normal formation of such circuits ^11, 12^. Evidence for this comes from studies in which whisker clipping ^13^, transection of the whisker afferent nerves ^14^, and manipulation of extrinsic serotonin levels ^15^ result in altered or absent intracortical projections from the barrel cortex.

One mechanism via which extrinsic inputs could promote the formation of appropriate cortico-cortical connections is patterned spontaneous activity that correlates depolarizations among disparate brain areas, allowing synaptic stabilization via Hebbian processes ^11, 12, 16, 17, 18^. Such mechanisms have been well described in visual systems ^19^ but evidence for them is weak in other systems. The large majority of studies of spontaneous activity patterns in early life have examined activity within a single sensory cortex ^20, 21, 22, 23, 24^. Studies using functional magnetic resonance imaging (fMRI) in very young human infants have shown networks of cortical areas synchronized by very slow oscillations of the BOLD signal ^25^, but how these networks relate to functional cortical networks at this age is not known. Others have examined the integration of sensory cortices into wider networks of cortical areas ^20, 26, 27^, but to date it is not known if these maturating networks are reflected in spontaneous cortical activity. If they are, it would provide a worthwhile target for further examination of the roles of extrinsic factors in the development of cortico-cortical networks.

In this study we address this issue by using voltage-sensitive dyes to provide signals of cortical activity with high spatial and temporal resolution ^28^ over a large region of the cortex spanning both medial cortical areas and the barrel cortex from the *in vivo* rat brain of young (P5-P12) pups. Our imaging was conducted under light anesthetic (0.25-0.75% isoflurane). We found that stimulation of the whisker in P5-6 pups evoked activity in a focused and local region of the barrel cortex. Stimulating the whisker in P12 pups resulted in activation of medial motor areas in addition to the barrel cortex. In the intervening ages (P8-P10) the medial activation was variable. Nevertheless, we found that spontaneous activity in the barrel cortex was correlated with cortical activity in the putative motor cortex, regardless of whether whisker stimulation evoked activity in these regions. These results suggest that spontaneous cortical activity may be a factor in the formation of mature cortico-cortical connections.

## Results

### Spatial and temporal properties of whisker-evoked activity change with age

Montages in Figure 1A show the cortical pattern of the whisker responses to contralateral whisker stimulation at five latencies in three pups, (P5, P8, P12; see also supplementary video 1-3). The latency of the VSD response decreased with age, from 83 ± 15 ms at P5 to 35 ± 5 ms at P12 (one-way ANOVA, p = 0.018). There was also a significant effect of age on time above 50% of the maximum response, which decreased from 363 ± 31 ms at P5 to 104 ± 24.2 ms at P12 (one-way ANOVA, p = 4.0 × 10^-4^). The mean area of cortex activated (see *Methods* for calculation details) increased with age from 0.40 ± 0.13 mm^2^ at P5 to 5.5 ± 1.0 mm^2^ at P12 (one-way ANOVA, p = 0.0012). These results are shown in Supplementary Figure 1. Also shown on Supplementary Figure 1 are the responses evoked by ipsilateral whisker stimulation. There was a significant effect of age on the maximum ipsilateral response (one-way ANOVA, p = 0.0022).

**Figure 1.**
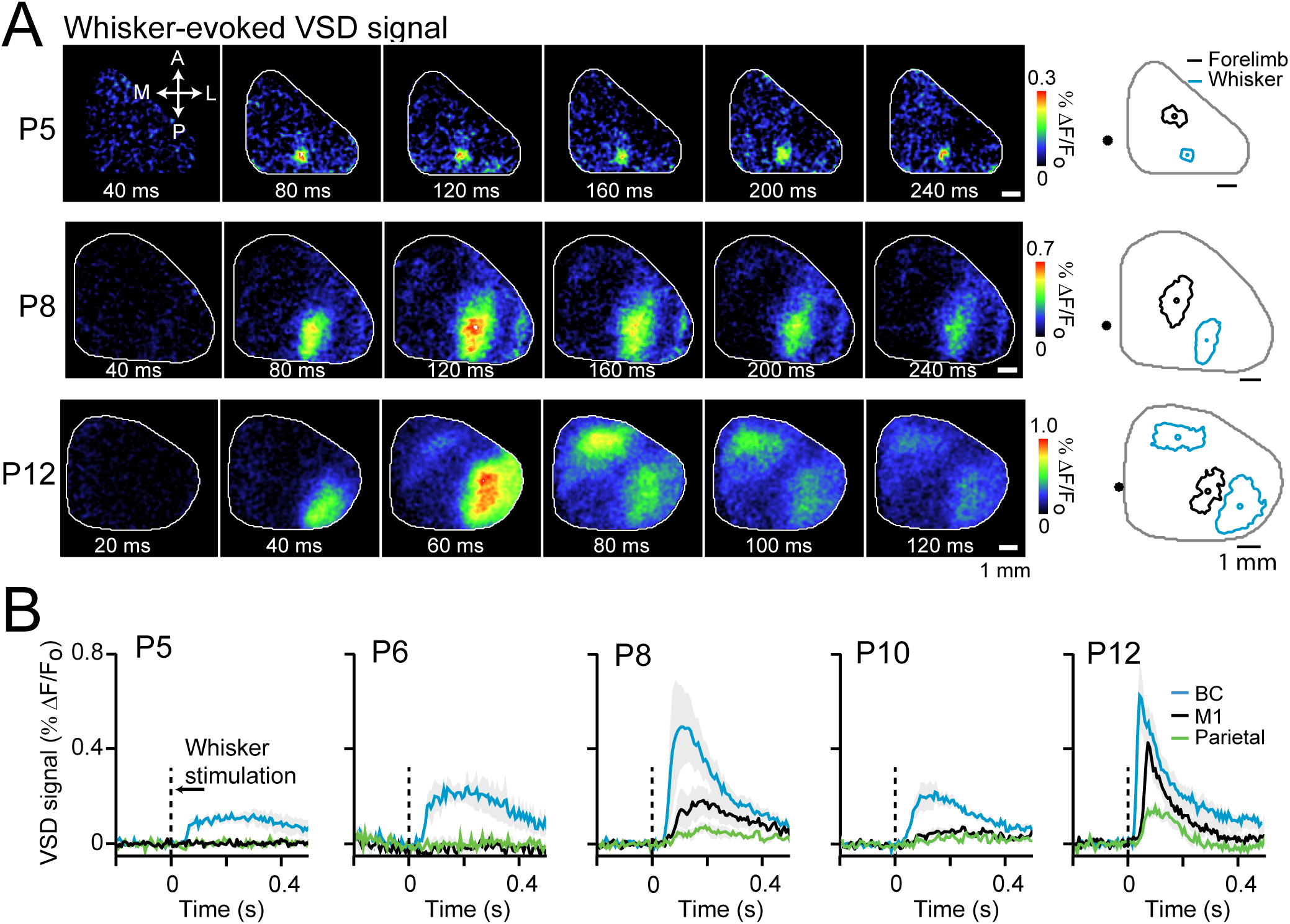
Voltage-sensitive dye imaging reveals patterns of cortical activation following sensory stimulation. **A)** Images showing patterns of cortical activation following whisker stimulation. First panel of P5 pup shows anterior (A), posterior (P), medial (M) and lateral (L) directions. Note different time intervals and scale bar different ages. At right, outlines of cortical regions reaching 60% of maximal VSD signal. Total imaged area outlined in gray. **B)** Mean signals, by age (P5, n = 4; P6, n = 5; P8, n = 5; P10, n = 4; P12, n = 7), collected from BC, M1, and parietal points following whisker stimulation. Dashed line represents time of whisker stimulation.

### Emergence of whisker-evoked activity in motor cortex

A prominent feature of the P12 cortical activation is the presence of an independent medial area of activity following stimulation of the whisker, similar to as described in the adult rodent ^4, 29, 30^. We describe this region as motor cortex based on this response and its location; we did not assess whether stimulation elicited movement responses. The outlines of these regions, along with the region activated by stimulating the forelimb, are shown in the left panels of Figure 1A.

To allow for a comparison of M1 activation following whisker stimulation across ages, we anatomically defined a region from which to collect a signal in this region in the absence of any detected depolarization during stimulation. We also defined a point directly medial to the barrel cortex distinct from the putative M1, for comparison (see *Methods* and Supplementary Figure 1 for details). We describe this latter point simply as *parietal* to differentiate it from the M1 point, within the frontal cortex. Figure 1B shows the mean VSD signals collected from these points following stimulation. Responses in M1 were absent at the earliest ages examined (P5 and P6), variable at P8 and P10 (we detected responses in 5/6 P8 pups and 2/4 P10 pups) and present in all pups at P12 (see also supplementary video 1-3).

To examine whether these responses were due to a second locus of activity and not simply a uniform spread of activation, we normalized the peak M1 response and the peak parietal response to the peak BC response. These ratios are shown in Supplemental Figure 1G for each age group. The normalized M1 response increased with age from 0.29 ± 0.025 at P5 to 0.63 ± 0.024 at P12 (one-way ANOVA, p = 0.0019). In contrast, the normalized parietal response did not change with age, being 0.32 ± 0.05 at P5 and 0.27 ± 0.038 at P12 (one-way ANOVA, p = 0.23). Two-way ANOVA of both sets of ratios across ages revealed an effect of age (p = 0.0128) and of cortical location (p = 7.0 × 10^-4^). Furthermore, the normalized M1 response was significantly correlated with the area of cortex activated following whisker stimulation (r = 0.61, p = 8.3 × 10^-4^) as well as the latency of activation following whisker stimulation (r = -0.48, p = 0.012). Importantly, the normalized parietal response was not significantly correlated with either the area of activation (p = 0.26) or the latency of activation (p = 0.48) following whisker stimulation. We interpret these results as evidence that the signal recorded from the medial anterior (presumptive motor) cortex is a distinct feature of maturation of the whisker sensorimotor system and not due to a general increase in the area of activation spread ^4, 29, 31^.

### Spontaneous cortical activity preferentially synchronizes motor and barrel cortices

Supplementary Figure 2 shows example sequences of spontaneous cortical activity from two pups of P6, and P12 ages, along with power and frequency properties of the VSD signals across all pups. We used the wide imaging area to examine the correlation of activity between the barrel and motor cortices during spontaneous activity. We observed that these regions could be active together spontaneously, even in those P5-6 pups in which whisker stimulation activated only the barrel cortex. Figure 2 shows three examples of this synchronous activation from three pups ranging from P5-P12 (see also supplementary video 1-3). Images on the left show sequential images of spontaneous brain activity, while the image on the right shows the sensory evoked pattern of activity (mean image calculated from onset to peak of activity). In these P5-P8 examples, the barrel cortex and the presumptive motor cortex are active together during spontaneous bursts, while sensory stimulation activated only the barrel cortex.

**Figure 2.**
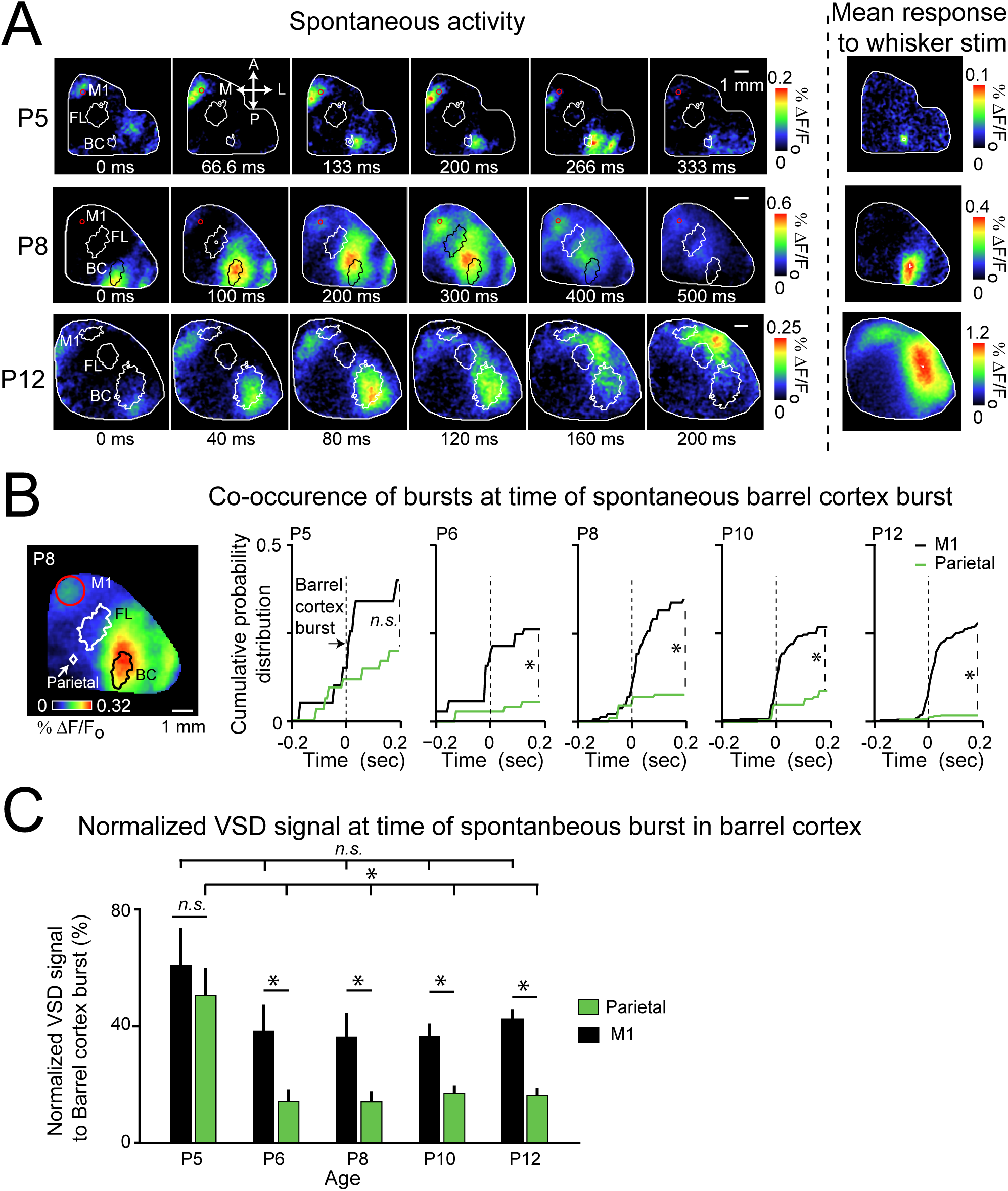
Synchronous activation of primary sensory cortex and motor cortex can exist during spontaneous activity even if absent during sensory stimulation. **A)** Images show example sequences in which barrel cortex (outlined) is activated synchronously with motor cortex. Forelimb region outlined for reference. Filled circle shows estimated M1 region, except in P12 example in which M1 activity was detected during sensory stimulation. Images at right of dashed line show activity image averaged from time of onset to time of peak of VSD signal following whisker stimulation. **B)** Mean image at time of burst in barrel cortex from one P8 pups. Putative M1 shown by red circle. At right, mean cumulative probability distribution of bursts in M1 (black) and parietal cortex (green) relative to burst in barrel cortex at different ages. Dashed line represents barrel cortex burst. **C)** VSD signal at time of barrel cortex burst (normalized to barrel cortex signal) in parietal (green) or motor (black) cortex. In this and all subsequent bar graphs, bars shown standard error and *\** indicates significant difference with exact value shown in text.

During whisker stimulation, activation of motor cortex follows activation of barrel cortex with a delay of 33.4 ± 5.4 ms at P12 (Fig 1A, B). To determine if this delay also exists during spontaneous activation of motor cortex, we calculated cumulative probability curves for the incidence of motor cortex bursts, relative to the timing of barrel cortex bursts (Figure 2B). This analysis was performed only at times where there was no activity in the forelimb cortex, which lies between the motor and barrel cortices. This ensured that spreading activity from the forelimb cortex would not be counted as co-incident bursts in the motor and barrel cortex. The occurrence of bursts in the motor cortex increases sharply at the time of bursts in the barrel cortices. This increase is absent in the parietal cortex. The overall difference in occurrence of bursts within 200 ms before or after a barrel cortex burst was significantly different in across all ages except P5 pups. (one-tail paired t-test of cumulative probability from 200 ms prior to 200 ms post barrel burst; P5, p = 0.20; P6, p = 0.029; P8, p = 0.011; P10, p = 0.035; P12, p = 6.4 × 10^-4^). Similarly, the normalized VSD signal in the motor cortex was higher than the normalized VSD signal in the parietal cortex at the time barrel cortex bursts in all ages, and significantly so in all ages except P5 (one-tailed paired t-test, P5, p = 0.11; P6, p = 0.046; P8, p = 0.034; P10, p = 0.0040; P12, p = 3.1 × 10^-4^; Figure 2C).

To examine this synchrony more closely, we generated correlation maps based on a seed pixel within the center of a distinct cortical area that showed the correlation of activity between each imaged point of cortex and a seed pixel during a 6-8 min period, as described in *Methods*. We generated such maps from both spontaneous and whisker-evoked activity, which allowed us to make direct comparisons between the degree of correlation between cortical points during spontaneous activity vs. during sensory-evoked activity.

Figure 3A shows example correlation maps from three pups. In all cases, the seed pixel is in the barrel cortex. In the three examples, there is a region of high correlation in the motor cortex in the maps generated from spontaneous activity. Note there is also a region of high correlation on the lateral edge of the imaged region, likely representing the whisker secondary sensory region ^29, 32^. The correlation maps generated from sensory-evoked activity, in contrast, has no region of higher correlation in the motor region in the P6 animal, a small weak region in the P8 animal, and a large robust region in the P12 animal (note the different scaling between the sensory-evoked and spontaneous maps). This is consistent with the emergence of M1 responses following whisker stimulation as shown in Figure 1. To control for spurious correlations, we calculated cross-correlation curves of VSD signal from motor and barrel cortex delayed by 2 seconds. Figure 3B shows summary mean cross-correlograms of simultaneous and delayed VSD signals. Note the loss of correlation with delay.

**Figure 3.**
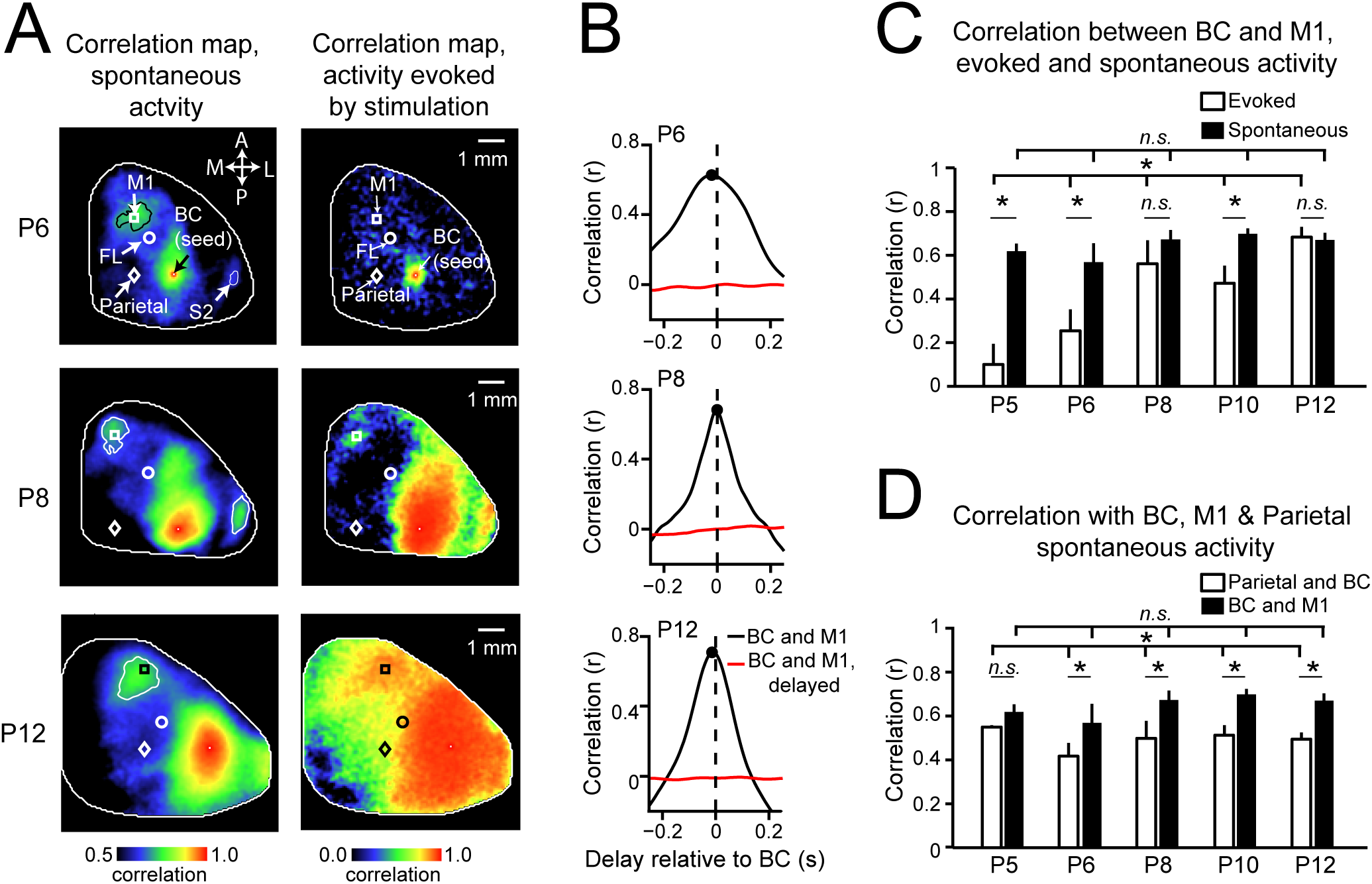
Activity in barrel and motor cortices is highly correlated during spontaneous activity, regardless of correlation during sensory activity. **A)** Maps showing correlation of activity at each point on imaged cortex with barrel cortex (seed pixel) during spontaneous activity (left) or activity following whisker stimulation (right). Forelimb (circle), M1 (square), and parietal (diamond) points shown for reference. Discrete islands of high correlation during spontaneous activity are outlined for emphasis. **B)** Mean cross-correlation of VSD signal in barrel and motor cortices, from all animals at three ages. Black, no delay; red, delay of two seconds. Black dot shows time of maximum correlation. **C)** Mean correlations between barrel and motor cortices during sensory evoked activity (white bars) and spontaneous activity (black bars). **D)** Correlations during spontaneous activity between parietal and BC (white) and M1 and BC (black).

Figure 3C compares the mean correlation across ages. The mean correlation between barrel and motor cortices during whisker evoked activity increased with age (0.010 ± 0.094 at P5, 0.69 ± 0.047 at P12; one-way ANOVA, p = 0.0015), while the correlation during spontaneous activity did not (0.62 ± 0.035 at P5, 0.67 ± 0.032 at P12; one-way ANOVA, p = 0.54). Results of paired one-tailed t-test comparing the barrel-M1 correlation values following sensory stimulation to those during spontaneous activity at each age were: P5, p = 0.0057; P6, p = 0.0069; P8, p = 0.24, P10, p = 0.017; P12, p = 0.69). Thus in three ages (P5, P6, P10) barrel and motor cortices were significantly more correlated during spontaneous activity than following whisker stimulation.

To examine the specificity of the relationship between barrel cortex and motor cortex, we compared these correlations to those between barrel cortex and the parietal point, as shown in Figure 3D. The correlation between barrel and motor cortices was significantly higher at all ages beginning at P6 (one tailed paired t-test, P5, p = 0.0645; P6, p = 0.0069; P8, p = 0.024; P10, p = 0.013; P12, p = 4.87 × 10^-4^), providing evidence that activity in the barrel cortex was not simply correlated indiscriminately with all cortical regions, but preferentially with M1.

### Spontaneous bursts of activity in barrel cortex associated with co-incident bursts in motor cortex

We considered two sources for the co-occurrence of bursts in barrel and motor cortex during spontaneous activity; subcortical areas, such as the thalamus, that coordinated related brain regions during spontaneous activity ^21, 23^; and intracortical connections that were inactive during sensory stimulation. To distinguish between these possibilities, we looked for evidence of intracortical connectivity at different ages, using intracortical microstimulation (ICMS) and anatomical tracing.

First, we used intracortical microstimulation to directly excite the barrel cortex at P5-6 and P12 pups (Figure 4). Whisker-evoked VSD response was used to identify the position of barrel cortex. Trains of electrical pulse were delivered into the barrel cortex using a tungsten electrode. ICMS stimulation evoked a response with short latency in the barrel cortex of P5-6 pups (n=3), which spread locally within the barrel cortex around the stimulation electrode (activation radius of 0.45 ± 0.12 mm; Figure 4A). The spatial pattern of the activity evoked by ISMS is similar to that induced by whisker stimulation, indicating that there is no functional connection between sensory and motor cortex at P5-6. ICMS stimulation at P12 pups, however activated the motor cortex, consistent with previous studies of the mature barrel cortex ^4, 29, 30^. Note that compared with the VSD activity spread in younger pups, the activity spread around the ICMS electrode in P12 pups was much larger after cortical response onset (0.45 ± 0.12 mm at P5-6 compared with 4.69 ± 0.87 mm at P12; one-way ANOVA, p = 0.0021).

**Figure 4.**
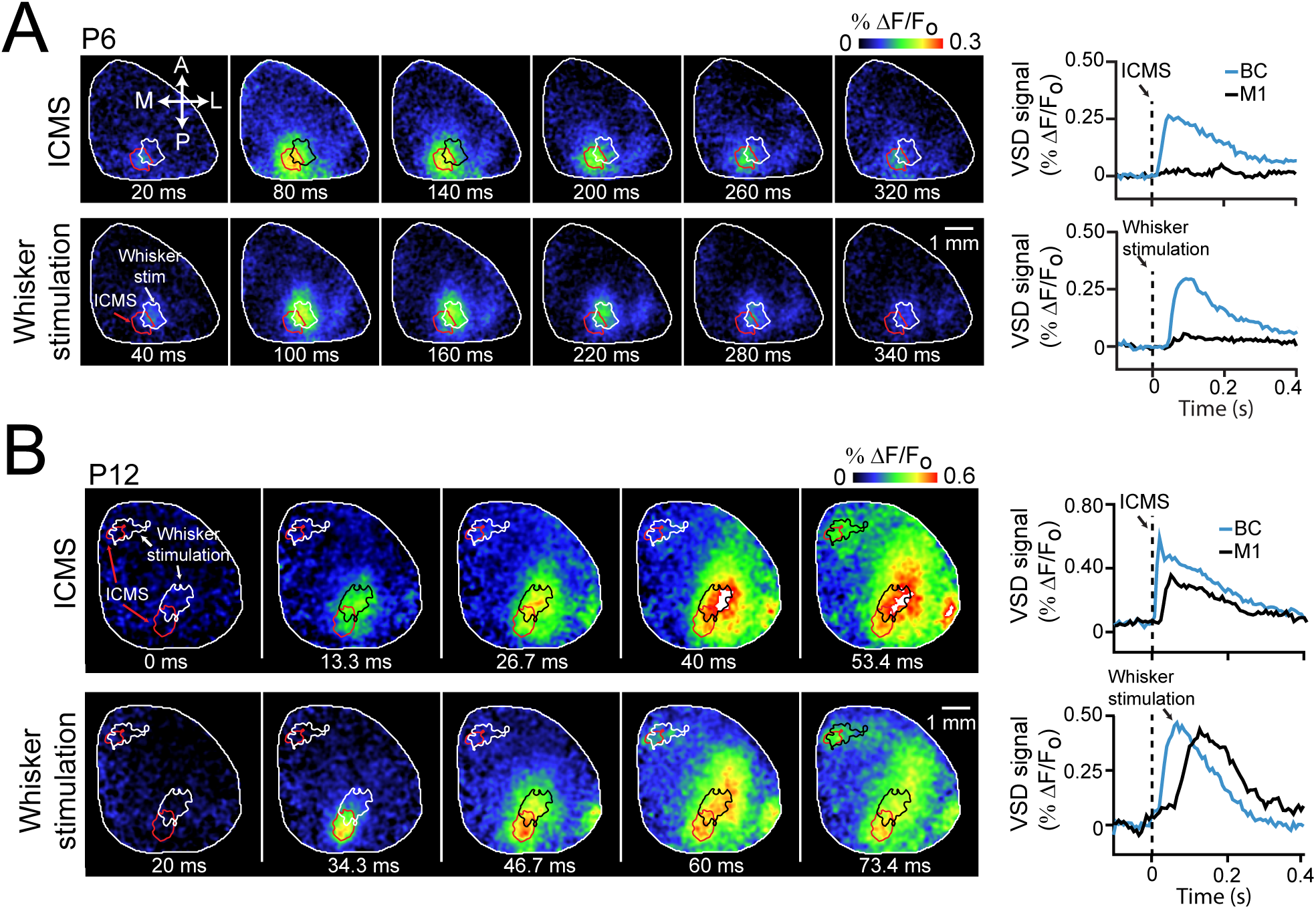
Intra-cortical micro-stimulation (ICMS) confirms the lack of functional connectivity between barrel and motor cortices at earlier ages. **A)** Left, representative montages of VSD signals following ICMS (upper) and whisker stimulation (lower). On both montages, cortical region activated by ICMS (red) and stimulation (white or black) is shown. Note the overlap of these activation regions. Right, average cortical responses from P5-P6 pups measured from barrel (blue) and motor (black) cortices following ICMS (upper) and whisker stimulation (lower). **B)** As in B, but in P12 pups. Note the robust activation of motor cortex following activation of sensory barrel cortex either by whisker stimulation or ICMS. Activation time differs due to direct cortical stimulation vs. ascending sensory stimulation.

In the second set of experiments, we specifically investigated whether neurons in the mouse barrel column project to M1 at different ages (Supplementary Figure 4). P4 and P10 pups (n=3 each group) were given a unilateral microinjection of the retrograde fluorescent tracer Fast blue ^33^ into the motor cortex (see *Methods*). For all animals analyzed (n = 6), the injection site spanned all cortical layers and spread ∼0.4-0.5 mm in diameter. At both P6 and P12 pups, fast blue labeling was evident within the homotopic region of the contralateral motor cortex (Supplementary Figure 4A, B), however the relative number of labeled neuron within barrel cortex differed significantly (P6/P12 ratio, 0.08 ± 0.01; p = 0.004). Connectivity was layer specific at P6. Compared with neurons in superficial layers of barrel cortex, more neurons were labeled in infragranular cortical layers (superficial/infragranular ratio, 0.09 ± 0.01; p = 0. 3 × 10^-4^). Our anatomical data provide evidence for a strong and direct glutamatergic connection from barrel cortex to motor cortex at P12 and not in P6.

## Discussion

Functional connectivity between sensory and motor cortices is a prominent feature of cortical circuits related to the rodent whisker system ^4^. In this study, we used voltage-sensitive dye imaging to examine the development of this connectivity in rat pups from the ages of P5-P12. Our main findings were that spontaneous activity in the barrel cortex is associated with activity in the motor cortex and that this association precedes the activation of the motor cortex following whisker stimulation.

### Earlier studies of the development of the whisker sensorimotor system

Two parallel threads of research intersect with the results presented here; studies that examine the development of cortical networks that respond to and process sensory stimuli; and studies that examine the developmental changes of spontaneous cortical activity in these same brain regions. Regarding the former stream of research, our results are in agreement with a series of previous studies showing that functional thalamocortical synapses form shortly after birth ^12, 21, 23, 27, 34, 35, 36, 37, 38^, as well as a recent study examining the maturation of EEG responses across a large expanse of cortex following whisker stimulation ^26^. Using intracortical electrodes, this study found weak activity in M1 emerged between P7 and P10, with much stronger activity detectable by epicranial electrodes emerging at P13. Similarly, we found activity in M1 following whisker stimulation in some animals at P8 and P10, with stronger activation by P12 (Figure 1).

With regards to spontaneous patterns of activity in the developing cortex, there is a rich description of activity in the somatosensory cortices ^39^ with a relative paucity of data regarding activity in other regions. The dominant form of activity in the sensory cortices is the spindle burst, a burst of ∼20 Hz oscillations lasting for 200-400 ms that synchronizes activity within the somatosenory or visual cortex ^12, 22, 40, 41, 42^. These fast oscillations are nested in slower, 1-4 Hz waves ^43^. At about P8, discrete bursts are replaced with continuous rhythmic activity ^44, 45^ and similarly, we found significant increasing activity in the 1-4 Hz band of frequency power first apparent at P8 (Supplementary Figure 2).

Relatively little is known about developmental activity in non-sensory regions. In one of the first studies of such activity, Seelke et al.,^46^ found a variety of patterns including long-lasting (2-5 s) slow transients, shorter discrete biphasic events, and bursts of gamma activity along the midline from occipital, parietal, and frontal (approximately what we describe as motor) cortices. The authors concluded in their discussion that, “…very little is currently known about the development of cortical activity outside primary sensory areas…”

### Synchronized activity in barrel and motor cortex during spontaneous activity precedes functional cortico-cortical connections

VSD imaging allows the collection of activity from both sensory and non-sensory areas simultaneously, making it especially useful to examine the correlation of activity between disparate regions. We found that activity in motor and sensory regions of cortex could be synchronized during spontaneous activity in the absence of any functional connections revealed by stimulation. Could there be activation of the motor cortex at these youngest ages that we failed to detect? Our results, combined with those of other studies, make it unlikely. We found that neither whisker stimulation nor direct barrel cortex stimulation activated the motor cortex at P5/6 – other studies have resulted in similar timelines for the development of responses in the motor cortex ^26^. Furthermore, inputs to the motor cortex arrive mainly from layer V of the barrel cortex ^47^ which in turn receives inputs primarily from layers 2/3 of the barrel cortex ^48^. Synaptic inputs to layer 2/3 of the barrel cortex are not mature until P12, and whisker stimulation does not reliably generate action potentials in cells of this layer until this time ^49^.

Where then could the synchronization we observed in spontaneous activity arise? The subplate, a transient structure present during development ^50^, is a likely source. Subplate neurons form bidirectional glutamatergic and GABAergic connections with both the thalamus and the developing cortex ^51, 52^ in some cases spanning wide expanses of cortex ^50^. It is thought to contribute to the synchronized fluctuations of BOLD signal observed in human infants. The subplate is essential for the generation of spindle bursts ^53^ via an amplification and processing of sensory signals such as muscle twitches or retinal waves generated in the periphery ^54^.

The thalamus is another possible source of synchronization during spontaneous activity as projections to both the motor and sensory ^37, 55, 56^ cortices are present at birth, and simultaneous recordings from the thalamus and the barrel cortex show tight correlation between the two ^21, 23^. Interestingly, temporal properties of spontaneous thalamic activity were different from those that followed whisker stimulation, as was the delay between the thalamic burst and the cortical burst ^23^. This suggests that spontaneous bursts of activity in the developing barrel cortex do not strictly reflect activation of ascending sensory pathways. Similarly, spindle bursts persist in the absence of peripheral inputs (via pharmacological transection of the spinal cord ^22^, inactivation of the whisker pad ^42^ or silencing of the retina ^40, 57^, which indicates activity intrinsic to the subplate or thalamus is an additional mechanism via which spindle bursts can arise. These different mechanisms could account for our observation that barrel and motor cortices could be activated simultaneously during spontaneous activity even when sensory stimulation activated only the barrel cortex.

### Possible functional role for synchronized activity in cortical networks

Widespread connectivity is an essential feature of the mammalian brain ^58^ and in the rodent, electrical recordings ^59^, imaging ^29, 31^ and optogenetics ^7^ have revealed functional interactions between discrete regions of cortex. Nevertheless we have only limited understanding of the development of the cortico-cortical connections that underlie this connectivity, especially in comparison to our understanding of the development of thalamocoritcal afferents and cortical patterning ^37, 60^.

A particularly important functional connection exists between S1 and M1 of the mammalian cortex, and spontaneous activity is uniquely suited to promote the maturation of this connection. Spontaneous activity in the developing cortex frequently results from involuntary muscle twitches of the limbs and whisker that occur during sleep ^39, 61, 62^. These twitches activate both sensory and proprioceptive receptors, which transmit feedback to S1 and M1, respectively ^24, 62^. In this way, twitch related feedback, amplified and transmitted by the subplate and thalamus can co-activate sensory and motor regions of the cortex. In contrast, during voluntary movements feedback from proprioceptive fibers is gated, limiting the coactivation of S1 and M1 ^27, 63^.

Mechanistically, it is known that axonal arbors are highly mobile during synaptogenesis, forming transient synapses with many possible post-synaptic target cells 64. Detection of correlated activity in pre- and post-synaptic cells via NMDA receptors and voltage-gated calcium channels prompts synaptic maturation ^65, 66^ which in turn stabilizes the parent branch via calcium-dependent mechanisms similar to those seen in the adult ^67^. Connections between neurons active together are strengthened, and those between unrelated neurons are lost ^68^. The co-activation of S1 and M1 following spontaneous twitches may provide coordinated spontaneous activity necessary for activity-dependent Hebbian plasticity.

## Materials and Methods

All procedures used in this study were conducted with the supervision and approval of the University of British Columbia or University of Lethbridge Animal Care Committee.

### Animal model and surgical procedures

We performed the experiments in this study using Sprague-Dawley rats of either sex, from ages of P5 to P12. We induced anesthesia using isoflurane (1.5-2%) mixed in oxygen. Following induction and throughout our imaging procedures, we maintained body temperature at 37 degrees C. Following subdural injection of local anesthetic, we removed the scalp and attached the skull to a custom built head-plate using dental cement. This headplate has embedded channels through which we passed warm water to maintain cortical temperature at 37 degrees, following the completion of preparatory surgery. We fixed the plate to a custom-built platform, and removed the skull and dura of the right hemisphere in a region extending approximately 2 mm posterior of bregma to 5 mm anterior of bregma, and immediately left of bregma to 7 mm lateral of bregma in rat pups of age P5-P12 (P5, n = 6; P6, n = 7 P8, n = 5; P10, n = 5; P12, n = 9). We clipped all whiskers on both sides except for the E2 whisker to allow for precise stimulation.

### VSD imaging

We dissolved the dye RH1692 (Optical Imaging) in HEPES-buffered saline, and applied it to the cortex exposed via the craniotomy described above. After allowing the dye to penetrate the cortex for 90 min, we removed it, washed the cortex with saline, and covered the craniotomy with 1.5% agarose, sealed with a glass coverslip. We transferred the steel plate with the attached rat pup to a mount fixed underneath our imaging system. This consists of a macroscope of front-to-front coupled video lens with a field of view of 8.4 square mm (65 ?m per pixel) with an associated 673-703 bandpass filter (Semrock, NY) as well as red LEDs (Luxeon K2, 627 nm) to excite the dye. We focused the camera ∼500 ?m under the surface of the cortex to reduce light distortion from vessels or other surface features. We collected images at 150 Hz using a digital camera (1M60 Pantera, Dalsa) and EPIX E8 frame grabber along with XCAP 3.8 software (EPIX). A second synchronized camera focused on the animal allowed us to monitor the pup for any large movements or jerks and adjust anesthesia as needed. The entire apparatus is located in a darkened enclosure isolated from external light and sound.

Images of cortical activity were collected under light isoflurane anesthesia (0.25-0.5%, P5-P8; 0.5-0.75% P10-P12). Under this level of anesthesia, myoclonic twitches were present in hindlimbs and tail (P5 = 0.091 +/-0.0064 Hz; P6 = 0.40 +/-0.011 Hz; P8 = 0.134 +/-0.0078 Hz; P10 = 0.031 +/-0.0021 Hz; P12 = 0.025 +/-0.0017 Hz). When collecting activity evoked by stimulation of the whisker or limbs, we illuminated the LEDs and began collecting images 100 ms later. Two hundred ms after the initiation of the camera, a single pulse of stimulation (∼200 ?m displacement over ∼10 ms) was given to the untrimmed E2 whisker or limb using a piezoelectric device (Q220-A4-203YB, Piezo Systems, Inc., Woburn, MA) and image collection was stopped 520 ms later (108 frames total). When stimulating the whisker, the stimulator was attached ∼ 1mm from the snout, resulting in a deflection of ∼10 degrees. When stimulating the forelimb, the stimulator was attached to the dorsal surface of the paw between the second and third digits. We also collected identical sets of images with no stimulation, to correct for time-dependant decreases in the VSD signal. For intracortical microstimulation experiments, trains of electrical pulse (50-100 µA, 1ms duration, 6 pulses, 5ms inter-pulse interval) were delivered into the E2 barrel cortex (depth of ∼400 µm) using a 0.1 M? unipolar tungsten electrode. E2 whisker sensory-evoked VSD response was used to identify the position of E2 barrel cortex. An overall sequence of images in response to stimulation was generated for each pup by taking the mean of 10-20 individual responses and filtering this mean with a Gaussian filter of 1 pixel standard deviation, using custom-script in ImageJ. We stimulated at most once every 10 s to prevent changes in response related to repetitive stimulation. The VSD responses were expressed as a percentage change relative to baseline VSD responses (ΔF/F0 × 100%) using ^®^Matlab (Mathworks, Natick, MA).

When collecting spontaneous cortical activity, we illuminated the LEDs and began collecting images 100 ms later. We collected images in 33.3 s (5000 frame) epochs. We filtered the images using a Butterworth zero phase-shift band-pass filter (0.5-6 Hz) and spatially filtered using a Gaussian filter of 1 pixel standard deviation. Change in fluorescence at time F was isolated from background signal by calculating (F – F_0_)/F_0_ where F_0_ is the average fluorescence. Finally, we reduced sources of noise by processing using principle-component analysis of each 5000 frame epoch and reconstructing the epoch using the first 40 principle components. We combined these epochs into sequences representing 6-10 min of activity for each pup and conducted our analysis on these aggregated image sets. To minimize movement artifacts, we eliminated sequences in which the pup made large jerks or movements as identified from the second synchronized camera.

### Tracing experiment

Rat pups were randomly divided into either the P4 or P10 group (3 rats per group), and selected to receive a primary motor cortex tracer injection. The rats were removed from their mothers and orally given ∼0.9 mg/kg of Meloxicam (0.5 mg/mL, Boegringer Ingelheim, Germany) 30 minutes prior to surgery. Five minutes prior to surgery, the rats were subcutaneously given 4 mg/kg of Lidocaine (20 mg/mL with 2% epinephrine, Western Drug Distribution Centre Ltd) over the incision site. Under isoflurane anesthesia the skull over the left hemisphere was exposed and a craniotomy was performed over the motor cortex (P4 AP 1.0 mm, ML 1.0 mm; P10 AP 2.0 mm, ML 2.0 mm). Fast Blue (2% w/v in ACSF) was injected (DV 0.4mm) in volumes of 350-400nL at 23 nL/s using a Nanoinject II (Drummond Scientific, Broomall, PA). The pups were sutured and returned to their mothers after they had regained sternal recumbency. Vanilla was applied to the mother’s nose to mask surgical smells. The rats were perfused two days post-surgery (P6 and P12, respectively). The brains were extracted and stored in 4% Paraformaldehyde (PFA) for 48h, then cryo-protected in 30% sucrose in 1x PBS until slicing. The brains were removed from sucrose and embedded in 3% agarose block for coronal vibratome sectioning at 40μm. Slices were mounted with Fluormount and subsequently imaged using a NanoZoomer 2.0-RS slide scanner (Hamamatsu, Shizuoka Prefecture, Japan) and 10x lens magnification.

### Data analysis

To determine the onset of cortical activity following stimulation of the whisker and limbs, we thresholded each image collected following stimulation at 60^th^ percentile. The first image on which a single area greater than 20 pixels^2^ (∼0.078 mm^2^) crossed this threshold was classified as the onset of activation. The center of this region was classified as the center of the sensory region being examined. We use the terms barrel cortex (BC) and and forelimb cortex (FL) to refer to the center of the regions activated by whisker and forelimb stimulation, respectively. To determine the area of activation, we calculated the mean image from the time of onset to the time of maximal VSD signal, and thresholded this mean image at the 60^th^ percentile. To determine if there was an independent activation in medial/motor regions of cortex following whisker stimulation, we examined the region of cortex anterior and medial to the detected center of forelimb S1. In this isolated region, we repeated the detection algorithm described above, searching for an isolated region of activated cortex larger than 20 pixels^2^ (∼0.078 mm^2^). If such an area was not found, we used a point 45 degrees anterior-medial to the detected forelimb region, at a distance of 75% of the distance between the detected forelimb and barrel regions. As this point represents the presumptive motor cortex, we termed this region primary motor cortex (*M1)*. As a comparison region of cortex, we used a point with the same medio-lateral position as the detected or calculated motor cortex, and the same anterior-posterior position as the detected barrel cortex. We refer to this point as simply *parietal* to differentiate it from the motor cortex point in the frontal cortex. These locations are summarized in Supplementary Figures 1C and D. We used these same points to analyze spontaneous cortical activity. For each region of interest (ROI) described above, we took the mean of the VSD signal from a 5x5 pixel region centered on the ROI.

To create correlation maps (Figure 3) we calculated the pair-wise correlation between the ROI VSD signal and the VSD signal from each other pixel. The value of the correlation was placed back into the position of the non-ROI pixel. We were interested to determine if there was a region of cortex in the anterior-medial region that was highly correlated with activity in the barrel cortex during spontaneous brain activity. In searching for such a region, we had two classes of pups – those in which the M1 point had been detected during whisker stimulation (see P12 pup in Figure 1 examples) and those the M1 point had been estimated, as no signal had been detected (see P8 and P5 pup in Figure 1 examples). In the case of the former, we collected the correlation value at the detected M1 point, and compared it to the parietal comparison point. In those pups in which M1 was estimated (not detected), we thresholded the correlation map anterior and medial to the FL S1 point at the 90^th^ percentile and took the center of this region. This strategy allowed us to find regions in the area of presumptive motor cortex that were correlated to the BC without any *a priori* knowledge of where exactly these regions would be.

To examine relationships between bursts in different cortical regions (Figure 2B), we set a threshold VSD signal above which we considered the region to be active. In pups age P5-P8, we determined the value of a threshold set at three times the standard deviation of the VSD signal. By age P10 and P12, cortical activity was nearly continuous (Supplementary Figure 2A) and accordingly the standard deviation was not appropriate for threholding. The values of three times the standard deviation for the P5-P8 pups was 0.10 +/-0.006 (BC), 0.11 +/-0.002 (M1), 0.10 +/-0.004 (parietal) and 0.10 +/-0.005 (FL). Thus, we used a threshold of 0.10 for the P10-P12 pups.

### Statistical analysis

We compared means between groups using one- or two-way anova as appropriate, with Tukey’s Honestly Significant Difference correction for multiple comparisons when comparing means of groups. We used paired one-way t-tests when comparing pairs of correlations. Error bars and ± ranges represent standard error. Correlation values were corrected using Fischer’s z-transformation before statistical analysis.

## Acknowledgments

This work was supported by a Natural Science and Engineering Council of Canada (NSERC) discovery grants #40352 to MHM and a Canadian Institutes of Health Research (CIHR) Operating Grant MOP-12675 and a Human Frontier Science Program grant to THM. MHM is a Campus Alberta for Innovation Program Chair, Alberta Alzheimer Research Program (MHM), and Alzheimer Society of Canada (MHM), CIHR Vanier Award and Michael Smith Foundation for Health Research predoctoral fellowship to DM. We thank Pumin Wang and Richard Kline for surgical assistance.

